# The Multifunctional Catalytic Hemoglobin from *Amphitrite ornata*: Protocols on Isolation, Taxonomic Identification, Protein Extraction, Purification, and Characterization

**DOI:** 10.1101/2024.11.20.624587

**Authors:** Anna L. Husted, Victoria R. Sutton, Hao V. Nguyen, Lauren A. Presnar, R. Kevin Blackburn, Joseph L. Staton, Stephen A. Borgianini, Edward L. D’Antonio

**Affiliations:** Department of Natural Sciences, University of South Carolina Beaufort, 1 University Boulevard, Bluffton, South Carolina 29909, USA; Department of Molecular and Structural Biochemistry, North Carolina State University, 120 W Broughton Drive, Raleigh, North Carolina 27607, USA

**Keywords:** marine benthic ecology, infauna, polychaete, allelochemical, halogenated aromatic compound, hemoglobin, dehaloperoxidase, molecular taxonomy, DNA barcoding, UV absorbance-based assay

## Abstract

The multifunctional catalytic hemoglobin from the terebellid polychaete *Amphitrite ornata*, also named dehaloperoxidase (*Ao*DHP), utilizes the typical oxygen transport function in addition to four observed activities involved in substrate oxidation. The multifunctional ability of *Ao*DHP is presently a rare observation, and there exists a limitation for how novel dehaloperoxidases can be identified from macrobenthic infauna. In order to discover more infaunal DHP-bearing candidates, we have devised a facilitated method for an accurate taxonomic identification that places visual and molecular taxonomic approaches in parallel. Traditional visual taxonomic species identification by the non-specialist, at least for *A. ornata*, or even for other marine worms, is a very difficult and time-consuming task since a large diversity is present and the method is restricted to adult worm specimens. The work herein aimed to describe a method that particularly simplifies the taxonomic identification of *A. ornata* through the assessment of its mitochondrial cytochrome c oxidase subunit I gene by employing the DNA barcoding technique. Furthermore, whole worm specimens of *A. ornata* were used to extract and purify *Ao*DHP followed by an H_2_O_2_-dependent peroxidase activity assay evaluation against substrate 2,4,6-trichlorophenol. *Ao*DHP isoenzyme A was also overexpressed as the recombinant protein in *Escherichia coli*, and its peroxidase activity parameters were compared to *Ao*DHP from the natural source. The activity assay assessment indicated a tight correlation for all Michaelis-Menten parameters evaluated. We conclude that the method described herein exhibits a streamlined approach to identify the polychaete *A. ornata*, which can be adopted by the non-specialist, and the full procedure is predicted to facilitate the discovery of novel dehaloperoxidases from other marine invertebrates.

## 1. Introduction

The polychaete worm *Amphitrite ornata* (Leidy, 1855) is found in marine soft-sediment habitats that contain significant volatile toxic halometabolites. Another polychaete worm, species *Rashgua lobatus* (Hartman, 1947) (with accepted name of *Notomastus lobatus*), and the enteropneust worm, species *Saccoglossus kowalevskii* (Agassiz, 1873), have been previously studied that can produce relatively high concentrations of toxic volatile bromometabolites [1-5]. Such compounds include 4-bromophenol, 2,4-dibromophenol, and 2,4,6-tribromophenol secreted by *N. lobatus* [1,5]; and 2,3,4-tribromopyrrole and 2,3,4-tribromopyrrole sulfamate produced by *S. kowalevskii* [2]. The primary purpose of these compounds is to act as negative cues to predation [6]. These hydrophobic allelochemicals are involved in bioaccumulation (worm tissue) and also the secretion into the surrounding sediments [1,7]. Fielman and colleagues have reported that other marine worms secrete added metabolites into the environment, presumably as negative cues as well [1]. Despite this chemical-based predator deterrence activity, *A. ornata* worms were not found to produce any detectable volatile toxic halometabolites, and were abundantly found inhabiting the same marine sedimentary environment as *N. lobatus* and *S. kowalevskii. Amphitrite ornata* worms are effectively protected by their coelomic hemoglobin from the harsh biogenic chemical toxicity, as they can utilize a H_2_O_2_-dependent peroxidase function from their respiratory pigment. This activity is enhanced and results in the oxidative dehalogenation of halometabolites to reduce them to lower cytotoxicity and increase the hydrophilicity of the resulting products [8]. Their hemoglobin was historically named dehaloperoxidase (*Ao*DHP) based on its notable peroxidase activity [8,9]. The enzymatic functions revealed by *Ao*DHP has permitted *A. ornata* to coexist in otherwise intolerable sediments produced by other benthic sea worms.

*Ao*DHP is a heme enzyme that was initially understood to perform solely in a bifunctional manner having peroxidase and oxygen transport/storage functions, and was classified as a dehaloperoxidase-hemoglobin [8-11]. However, three more functions were discovered in the past decade where the enzyme could also demonstrate oxidase [12], peroxygenase [12], and oxygenase [13] activities. Consequently, this unique five-function ability has redesignated *Ao*DHP as a multifunctional catalytic hemoglobin [14]. *Ao*DHP is also found in nature as two isoenzymes, *Ao*DHP-A and *Ao*DHP-B [15], and both of these forms have been studied biochemically for their structure and function [8,10,11,16,17]. Forms A and B differ by five amino acid residues, and biochemical studies have revealed some differences in their enzymatic performance; however, a side-by-side structural comparison between *Ao*DHP-A and *Ao*DHP-B was found to be very similar [8,14,17]. In solution, the quaternary structure of *Ao*DHP is monomeric and it has a molecular weight of 15.5 kDa [18,19]. The active site contains a heme prosthetic group where the iron ion is coordinated to an imidazole nitrogen of residue H89 (proximal histidine) [8]. In the presence of hydrogen peroxide, a common substrate such as 2,4,6-tribromophenol effectively undergoes a two-electron oxidation to ultimately form the product 2,6-dibromoquinone, in which a para-positioned bromine atom is displaced from the phenolic compound [11,19]. In general, given the importance of *A. ornata* sea worms (with regard to their *Ao*DHP production) notwithstanding, it has been quite challenging to accurately identify them, and nevertheless other worm species, by only visual taxonomy when the field research is conducted by a non-specialist. This challenge arises from finding juvenile worms that are difficult to characterize because they are not fully developed, thus there is a limitation in finding adult specimens throughout a given year [20,21]. There is also the problem of having an abundance of other marine worm taxa to try to distinguish from, which may add to the confusion [21,22]. The misidentification of any taxa would also result in profound negative consequences for research in the field [22]. In this work, we introduce a new method that removes these complexities in the isolation process and combines visual taxonomy (at a basic level) with molecular taxonomy that are both used in parallel. The overall goal is to streamline the processes of isolation and taxonomic identification but also keeping in line with the previous flowchart devised by Chen and colleagues on the purification of *Ao*DHP from *A. ornata* worms [19].

Herein, we present on an optimized set of protocols for (*i*) the isolation of *A. ornata* worms from a coastal benthic marine habitat in South Carolina using standard visual taxonomic procedures; (*ii*) the species-level identification of *A. ornata* by a molecular taxonomic approach involving the DNA barcoding of the mitochondrial cytochrome c oxidase subunit I (COI) gene; and (*iii*) the protein extraction, purification, and characterization of *A. ornata* dehaloperoxidase from the natural source (*Ao*DHP_NS_). The overall procedure was partly modeled from the work of Chen and co-workers [19]. Through characterization, a trypsin digestion followed by liquid chromatography with tandem mass spectrometry (LC-MS/MS) was performed and a comparison was made between *Ao*DHP_NS_ to that of *Ao*DHP overexpressed and purified as the recombinant protein from *Escherichia coli* (*Ao*DHP_REC_-A). Finally, a H_2_O_2_-dependent peroxidase assay comparison was made between *Ao*DHP_NS_ and *Ao*DHP_REC_-A. Kinetics for Michaelis-Menten parameters between *Ao*DHP_NS_ and *Ao*DHP_REC_-A revealed values to be in very close agreement.

## 2. Experimental Design

The outlined procedures described here are for the isolation and taxonomic identification of *A. ornata* sea worms, along with the extraction, purification, and kinetic characterization of *Ao*DHP_NS_ from *A. ornata* specimens. The overexpression (as carried out in *E. coli* host cells), purification, and kinetic characterization of *Ao*DHP_REC_-A is also described (**Figure 1**). The steps are presented in suitable detail with high-resolution figures included as accommodating visual aids.

**Figure 1.**
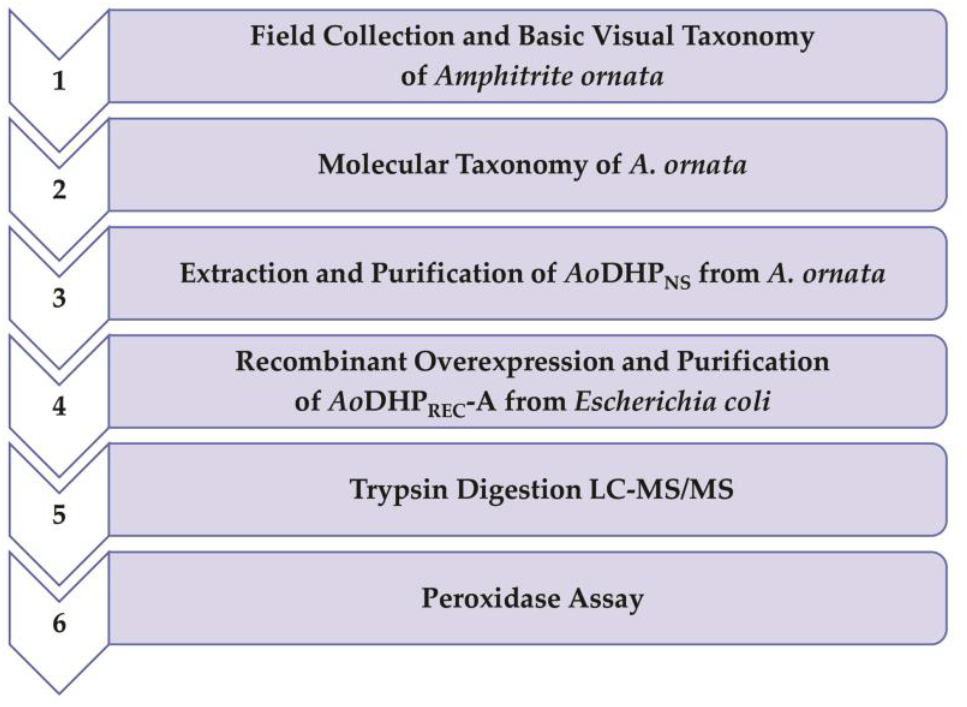
Methodological flowchart for the isolation and taxonomic identification of *A. ornata* marine worms, followed by the extraction, purification, and characterization of *Ao*DHP from the natural source and through recombinant overexpression.

### 2.1. Chemicals and Materials

1. 10x Tris/Glycine Running Buffer (Bio-Rad; Hercules, CA, USA; cat. no. 1610732).
2. 1kb DNA ladder (Promega; Madison, WI, USA; cat. no. G5711).
3. 2,4,6-Trichlorophenol (Sigma; St. Louis, MO, USA; cat. no. T55301).
4. 2,6-Dichloro-1,4-benzoquinone (Sigma; St. Louis, MO, USA; cat. no. 431982).
5. 2-Mercaptoethanol (Sigma; St. Louis, MO, USA; cat. no. M3148).
6. 2x Yeast Extract Tryptone (2xYT) Medium (Fisher Scientific; Waltham, MA, USA; cat. no. DF0440-17).
7. 4-(2-Hydroxyethyl)piperazine-1-ethanesulfonic acid (HEPES) (Sigma; St. Louis, MO, USA; cat. no. H3375).
8. 50x TAE Buffer (Fisher Scientific; Waltham, MA, USA; cat. no. BP13321).
9. Spectrum Spectra/Por 1 RC Dialysis Membrane Tubing 6-8 kDa (Fisher Scientific; Waltham, MA, USA; cat. no. 08-670C).
10. 6x Blue/Orange Loading Dye (Promega; Madison, WI, USA; cat. no. G1881).
11. Acetic acid (Sigma; St. Louis, MO, USA; cat. no. 695092).
12. Acetonitrile (Sigma; St. Louis, MO, USA; cat. no. 34851).
13. Agarose (Low EEO, Molecular Biology Grade) (Sigma; St. Louis, MO, USA; cat. no. A9539).
14. *Ao*COI-FORW Primer: 5’-CTCCATAAGATTACTAATTCG-3’ (Integrated DNA Technologies, Inc.; Coralville, IA, USA; cat. no. Custom Order).
15. *Ao*COI-REVR Primer: 5’-CTGATGGGTCAAAGAAAGAAG-3’ (Integrated DNA Technologies, Inc.; Coralville, IA, USA; cat. no. Custom Order)
16. Coomassie Brilliant Blue R-250 (Fisher Scientific; Waltham, MA, USA; cat. no. 04-821-616).
17. Buffer EB (Qiagen; Germantown, MD, USA; cat. no. 19086).
18. CM Sepharose Fast Flow Cation Exchange Media (Cytiva; Uppsala, Sweden; cat. no. 45-002-932).
19. Deoxyribonuclease I (DNase I) from bovine pancreas (Sigma; St. Louis, MO, USA; cat. no. DN25).
20. Dimethyl sulfoxide (DMSO) (Sigma; St. Louis, MO, USA; cat. no. 472301).
21. DNeasy Blood & Tissue Kit for DNA Isolation (Qiagen; Germantown, MD, USA; cat. no. 69504).
22. *E. coli* strain BL21(DE3) (New England Biolabs; Ipswich, MA, USA; cat. no. C2527H).
23. Ethidium bromide solution (10 mg/mL) (Sigma; St. Louis, MO, USA; cat. no. E1510).
24. Formic acid (Sigma; St. Louis, MO, USA; cat. no. 695076).
25. Hemin from porcine (Sigma; St. Louis, MO, USA; cat. no. 51280).
26. Hydrogen peroxide solution (30% wt.%) (Sigma; St. Louis, MO, USA; cat. no. 216763-100ML).
27. Hydrophilic 0.22 μM PVDF Filter (Fisher Scientific; Waltham, MA, USA; cat. no. SLGV033NS).
28. Imidazole (Sigma; St. Louis, MO, USA; cat. no. I2399).
29. Isopropanol (91%) (Fisher Scientific; Waltham, MA, USA; cat. no. 01-334-895).
30. Isopropanol (99.9%) (Sigma; St. Louis, MO, USA; cat. no. 34863).
31. Kanamycin sulfate (Fisher Scientific; Waltham, MA, USA; cat. no. BP906-5).
32. Laemmli (2x) (Bio-Rad; Hercules, CA, USA; cat. no. 161-0737).
33. Luria-Bertani (LB) broth, Miller (Fisher Scientific; Waltham, MA, USA; cat. no. BP1426-500).
34. Lysozyme (Fisher Scientific; Waltham, MA, USA; cat. no. PI89833).
35. Methanol (Fisher Scientific; Waltham, MA, USA; cat. no. A412-4).
36. Mini-Protean TGX 4-15% polyacrylamide gel (Bio-Rad; Hercules, CA, USA; cat. no. 456-1086).
37. Potassium chloride (Sigma; St. Louis, MO, USA; cat. no. P3911).
38. Potassium ferricyanide(III) (Sigma; St. Louis, MO, USA; cat. no. 702587).
39. Potassium phosphate dibasic anhydrous (Fisher Scientific; Waltham, MA, USA; cat. no. P288-500).
40. Potassium phosphate monobasic (Fisher Scientific; Waltham, MA, USA; cat. no. BP362-500).
41. Precision Plus Protein Dual Color Standards (Bio-Rad; Hercules, CA, USA; cat. no. 161-0374).
42. Protease Inhibitor Tablets (EDTA-free) (Fisher Scientific; Waltham, MA, USA; cat. no. A32965).
43. QIAquick Gel Extraction Kit (Qiagen; Germantown, MD, USA; cat. no. 28704).
44. Ribonuclease A (RNase A) from bovine pancreas (Sigma; St. Louis, MO, USA; cat. no. R4875).
45. Sodium chloride (Fisher Scientific; Waltham, MA, USA; cat. no. S271-500).
46. Sodium hydroxide (Sigma; St. Louis, MO, USA; cat. no. 221465).
47. Talon Co2+-NTA resin (Fisher Scientific; Waltham, MA, USA; cat. no. 89964).
48. *Taq* PCR Kit (New England Biolabs; Ipswich, MA, USA; cat. no. E5000S).
49. Tris base (Fisher Scientific; Waltham, MA, USA; cat. no. BP152-500).
50. Trypsin (Promega; Madison, WI, USA; cat. no. V5111).

### 2.2. Equipment

1. Quartz Cuvette Microcell (1.0-cm Pathlength) (Starna Cells; Atascadero, CA, USA; cat. no. 16.100-Q-10/Z15).
2. 5-mL Q-HP Chromatography Column (Cytiva; Uppsala, Sweden; cat. no. 17115401).
3. Agilent 8453 UV-visible Spectrophotometer (Agilent Technologies; Santa Clara, CA, USA; cat. no. G1103A).
4. ÄKTA Prime Plus FPLC System (GE Healthcare; Piscataway, NJ, USA; cat. no. 11001313).
5. Allegra X-30R Refrigerated Centrifuge (Beckman Coulter, Inc.; Brea, CA, USA; cat. no. B06320).
6. Amicon Ultra-15 Centrifugal Concentrator (MWCO: 10 kDa) (Millipore; Burlington, MA, USA; cat. no. UFC901024).
7. BMG VANTAstar F Microplate Reader (BMG LABTECH, Inc.; Cary, NC, USA).
8. Micro Pestle (United Scientific Supplies, Inc.; Libertyville, IL, USA; cat. no. 81441).
9. Peristaltic Pump P-1 (Cytiva; Uppsala, Sweden; cat. no. 18111091).
10. MyCycler Thermal Cycler (Bio-Rad; Hercules, CA, USA; cat. no. 170-9703).
11. QSonica Sonicator Q55 (QSonica, LLC; Newtown, CT, USA; cat. no. Q55).
12. ÄKTA 50-mL Superloop (GE Healthcare; Piscataway, NJ, USA; cat. no. 18111382).
13. UV-Star Microplate, 96-well, Clear, F-Bottom (Greiner Bio-One GmbH; Frickenhausen, Germany; cat. no. 655801).
14. Sterilmatic STM-E Steam Sterilizer (Market Forge Industries, Inc.; Everett, MA, USA; cat. no. STM-E).

## 3. Procedure

### 3.1. Amphitrite ornata Field Collection, Cleaning, and Cryopreservation

1. Obtain a scientific collection permit from the South Carolina Department of Natural Resources (SCDNR);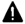 **CRITICAL STEP** Contact an agent or fill out an application online at the following webpage: https://www.dnr.sc.gov/.
2. Arrive to an oyster reef in a coastal marine environment of South Carolina (survey sites include sections on Hilton Head Island and Pritchards Island);
  a. Coastal marine field conditions: oyster reefs in an intertidal zone at low-tide with low-energy waves (**Figure 2**);
  b. High salinity;
  c. Outside temperature range: 70 – 90 °F;
  d. Months to find adult specimens: March – September;
3. Search for soft plough mud that is adjacent to a set of oysters (**Figure 2**);
4. Wear nitrile gloves;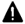 **CRITICAL STEP** To prevent contamination, one should wear gloves so as to not transfer any of their DNA from their hands onto the worm samples.
5. Dig approximately one foot under the oysters using a drain spade shovel;
6. Search for *A. ornata* worm specimens using basic visual taxonomic characteristics (**Figure 3a**) [23];
  a. Length of an adult worm is between 10 – 13 cm (4 – 5 inches) long;
  b. Color can be bright to dark red;
  c. One side contains spaghetti-like tentacles;
  d. Host of a U-shaped burrow;
  e. Symbionts associated with the ornate terebellid worm (*A. ornata*) include *Lepidametria commensalis* (*Lepidasthenia commensalis*) [24] and the tube pea crab (*Pinnixa chaetopterana) [25];*
7. Carefully place worm specimens into a plastic storage bag within an ice chest;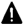 **CRITICAL STEP** Use new storage bags to prevent DNA contamination.
8. Return to the laboratory within three hrs;
9. Clean worm specimens from plough mud debris using a cold saline neutral buffer, such as 100 mM potassium phosphate (pH 7.0) in a clean plastic weigh boat/petri dish (**Figure 3b**);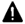 **CRITICAL STEP** Do not use distilled water during the cleaning process since there will be a deficiency in its salinity and the worm specimens will quickly perish thus compromising the *Ao*DHP_NS_ sample.
10. **OPTIONAL STEP** If an in-house *A. ornata* zoological specimen is desired, take one specimen and it is best to use 91% (v/v) isopropanol for additional debris removal, followed by its storage in fresh isopropanol solution. Note: Once the specimen has come into contact with the alcohol, it is no longer viable for any *Ao*DHP_NS_ extraction.
11. From the cleaned worm specimens (step no. 9), approximately 1.0 cm of the tail was removed by using a sterile scalpel knife;
12. The tissue of the worm specimen tail is carefully transferred into a sterile 1.5-mL Eppendorf tube that will be used for genomic DNA (gDNA) extraction, in the molecular taxonomic procedure (*vide infra*). Note: Label Eppendorf tubes appropriately to the corresponding whole worm tissue and cross-reference to the laboratory notebook.
13. The cut tail pieces are stored at -20 °C until the gDNA extraction protocol commences;
14. Individually, place whole worm specimens (dry and clean) into new plastic storage bags that are properly labeled by being cross-referenced to their corresponding cut tail pieces and to the laboratory notebook;
15. The whole worm specimens are stored at -80 °C for proper cryopreservation.

**Figure 2.**
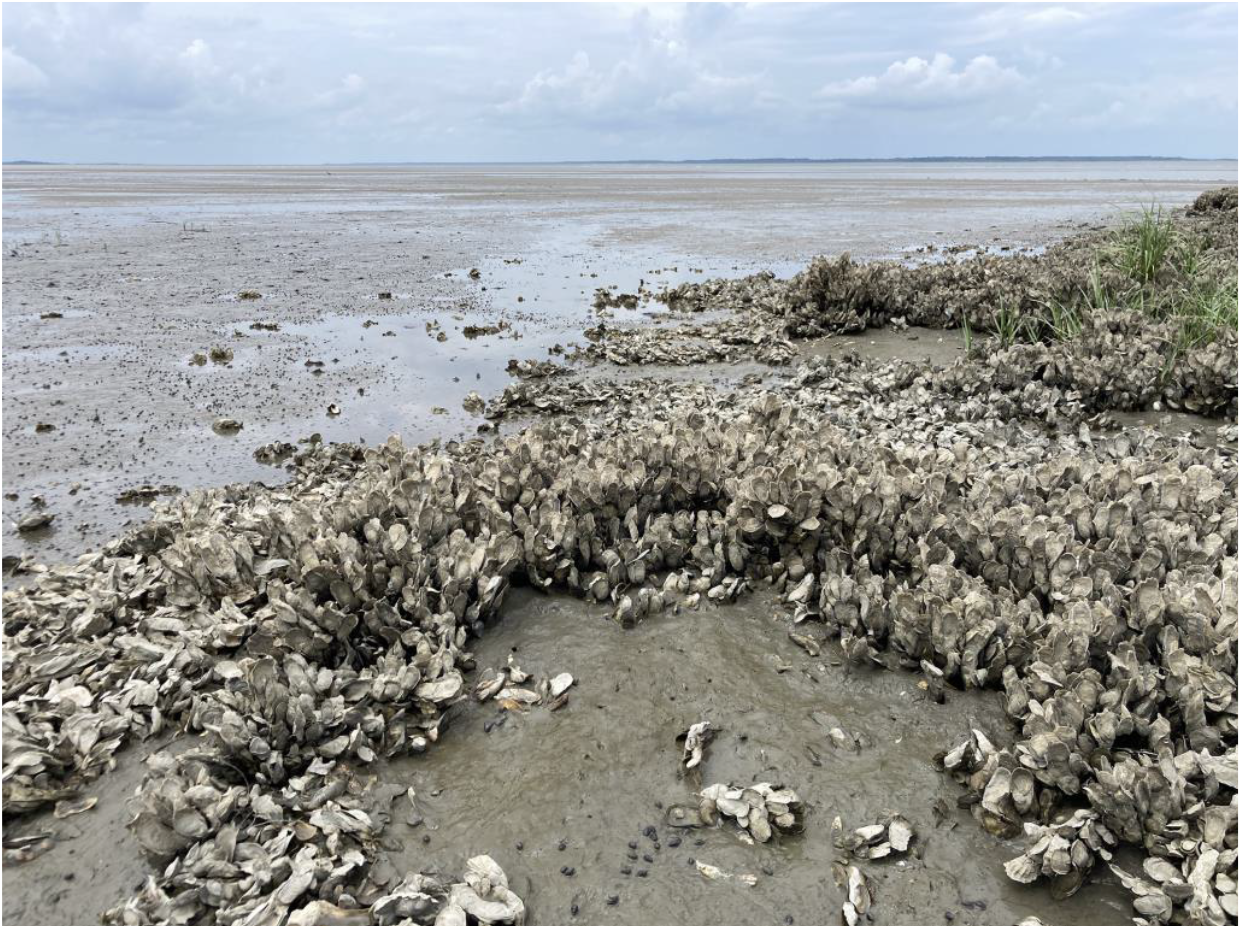
Field collection site for *A. ornata* marine worms at Hickory Forest Beach in Hilton Head Island, South Carolina, USA; coordinates: 32.2509536, -80.6974940. This private beach includes a variety of healthy oyster reefs in soft plough mud and high salinity, which are the ideal areas to find *A. ornata* in addition to other polychaetes. The image was taken from one of the oyster reefs in the intertidal zone during low tide.

**Figure 3.**
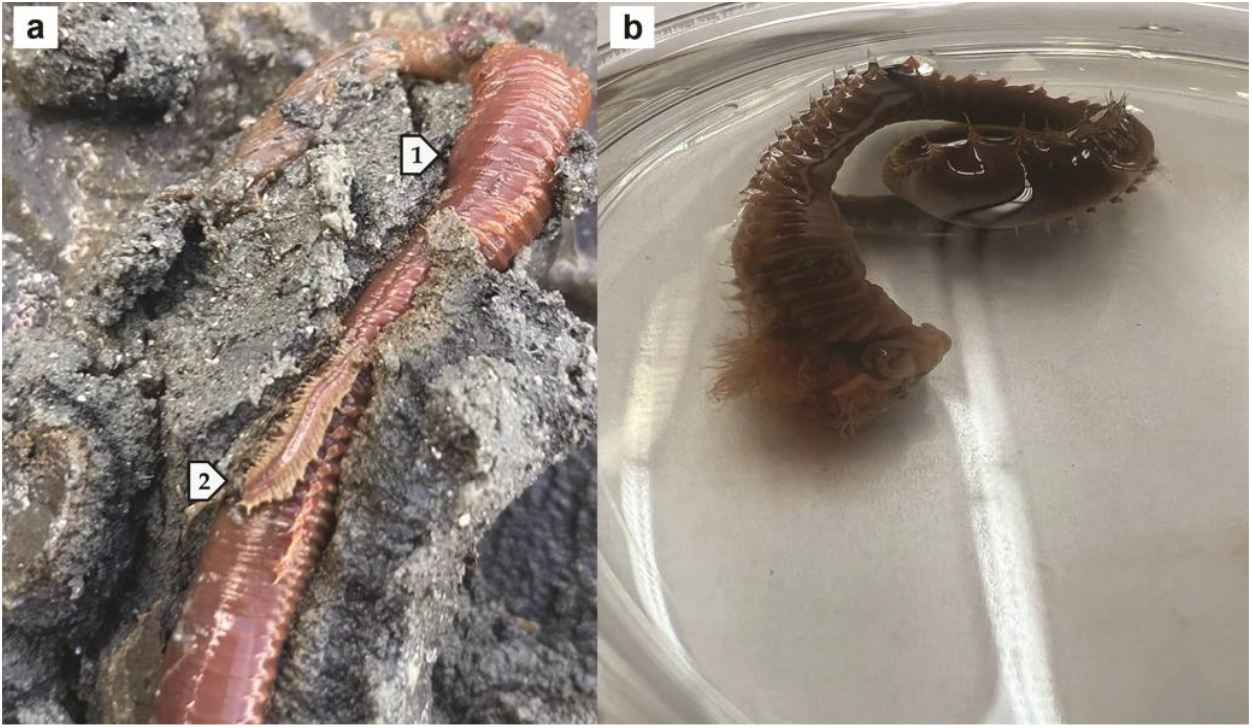
Field collection and cleaning of marine polychaete worms. (a) Excavated polychaetes from plough mud, including the host burrower (1) the ornate terebellid worm (*Amphitrite ornata*) (length: 7.5 cm x width: 0.6 cm) in the presence of one of its symbionts, (2) *Lepidametria commensalis* (length: 1.9 cm x width: 0.4 cm). (b) The *A. ornata* specimen was cleaned from mud debris in a neutral saline buffer, 100 mM potassium phosphate (pH 7.0).

### 3.2. Molecular Taxonomy

#### 3.2.1. Genomic DNA Extraction from A. ornata

1. Obtain the DNeasy Blood & Tissue Kit for DNA Isolation (Qiagen);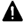**CRITICAL STEPS** Throughout this procedure, use sterilized pipette tips and 1.5-mL Eppendorf tubes in order to avoid DNA contamination. Use a refrigerated centrifuge set to a temperature of 4 °C for all centrifugation steps. All buffers indicated are from Qiagen.
2. For a given specimen, remove the 1.5-mL Eppendorf tube from the freezer containing the worm’s tail piece “tissue” (see step 13 in section 3.1) and thaw to room temperature for 15 min;
3. Add 180 µL of buffer ATL;
4. Grind the tissue with an autoclaved micro pestle in the 1.5-mL Eppendorf tube;
5. Add 20 µL of proteinase K (Qiagen) followed by vortexing;
6. Add 200 µL of buffer AL and continue to grind until the consistency of a paste is attained;
7. Incubate sample at 56 °C for 1.0 hr with vortexing every 15 min (note: the tissue should merge well into the liquid, and incubation may take a longer duration if the tissue samples are larger and not completely dissolved at the end of the 1.0 hr period);
8. Transfer the sample into a spin column;
9. Centrifuge for 1.0 min at 6,000 rpm;
10. Transfer the flow-through into a new spin column;
11. Add 500 µL of buffer AW1;
12. Centrifuge for 5.0 min at 6,000 rpm;
13. Discard the flow-through;
14. Add 500 µL of buffer AW2;
15. Centrifuge for 11 min at 13,000 rpm;
16. Discard the flow-through;
17. Transfer the spin column to a 1.5-mL Eppendorf tube;
18. Add 200 µL of buffer AE directly to the membrane of the spin column and allow it to soak for 1.0 min at room temperature;
19. Centrifuge for 2.0 min at 6,000 rpm;
20. **OPTIONAL STEP:** For a higher concentration of gDNA, add 200 µL of the flow-through (from step 19) back into the spin column and repeat steps beginning from step 11 (*vide supra*);
21. Store the flow-through at -20 °C and properly label the gDNA sample, which will be used in the PCR protocol.

#### 3.2.2. Polymerase Chain Reaction to Acquire the COI Amplicon of A. ornata

1. Obtain the *Taq* PCR Kit (New England Biolabs);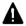**CRITICAL STEPS** Unless otherwise specified, all unique reagents in this procedure are provided in the *Taq* PCR kit. Sterilized pipette tips and tubes are required in this procedure in order to avoid DNA contamination.
2. Obtain the custom *A. ornata*-specific COI forward primer oligonucleotide DNA sequence, *Ao*COI-FORW: 5’-CTCCATAAGATTACTAATTCG-3’ and prepare a solution of 10.0 μM using buffer EB (Qiagen) as the diluent;
3. Obtain the custom *A. ornata*-specific COI reverse primer oligonucleotide DNA sequence, *Ao*COI-REVR: 5’-CTGATGGGTCAAAGAAAGAAG-3’ and prepare a solution of 10.0 μM using buffer EB (Qiagen) as the diluent;
4. Obtain a single PCR tube;
  a. Add 82.5 µL of nuclease free deionized water;
  b. Add 10.0 µL of 10x standard *Taq* buffer;
  c. Mix solution gently using a micropipette;
  d. Add 2.0 µL of 10 mM dNTPs;
  e. Add 2.0 µL of 10.0 µM forward primer (*Ao*COI-FORW);
  f. Add 2.0 µL of 10.0 µM reverse primer (*Ao*COI-REVR);
  g. Add 1.0 µL of template DNA after thawing to room temperature for 15 min (note: the template DNA in this case is the gDNA extracted from the *A. ornata* specimen, see step 21 in section 3.2.1);
  h. Add 0.5 µL of *Taq* polymerase;
  i. Mix gently with a micropipette;
5. Create a program in a PCR thermal cycler, as follows;
  a. Initial denaturation: 95 °C for 30 sec (1 cycle);
  b. Melt, Anneal, and Extension (40 cycles);
    i. Melt: 95 °C for 20 sec;
    ii. Anneal: 45 °C for 30 sec;
    iii. Extension: 68 °C for 60 sec;
  c. Final elongation: 68 °C for 5 min (1 cycle).
6. Place the PCR tube containing all reagents added from step 4.i. into the PCR thermal cycler and run the program from step 5;
7. Store PCR tube at 4 °C, which will be used in the agarose gel electrophoresis procedure.

#### 3.2.3. COI Amplicon Purification: Agarose Gel Electrophoresis and Gel Band Excision

1. Prepare 1000 mL of a 1x TAE solution by mixing 980 mL of deionized water with 20 mL of 50x TAE stock solution;
2. Cast a 1.2% (w/v) agarose gel;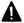**CRITICAL STEP** One must wear nitrile gloves during all procedural steps in this section in order to avoid exposure to ethidium bromide. Please refer to the MSDS of this chemical for information regarding safety and handling.
  a. Add dams into the gel pouring plate;
  b. Using a 250 mL Erlenmeyer flask (made of glass), mix 50 mL of 1x TAE solution with 0.600 g of agarose;
  c. Microwave the mixture for 45 sec;
  d. Swirl until most or all of the agarose is dissolved;
  e. Microwave again for an additional 30 sec and swirl again (note: the agarose should be fully dissolved);
  f. Pipet 10.0 µL of 10 mg/mL ethidium bromide (EtBr) solution into the agarose/TAE solution;
  g. Swirl gently;
  h. Immediately pour the agarose/TAE/EtBr solution into the gel pouring plate and install a 4-lane comb for the wells (notes: ensure that the comb does not touch the bottom of the pouring plate as this will break the gel; when pouring in the solution, work carefully as to not introduce bubbles);
  i. Allow the gel to cool for 20 – 30 min;
3. Prepare the DNA ladder;
  a. Add 40.0 µL of the 1 kb DNA ladder (Promega) into a clean 1.5-mL Eppendorf tube;
  b. Add 8.0 µL of blue/orange 6x gel loading dye (Promega);
  c. Mix gently with a micropipette;
4. Add 20.0 µL of blue/orange 6x gel loading dye (Promega) to a 100 µL PCR amplicon sample (see step 7 in section 3.2.2);
5. Cover the solidified gel with 1x TAE solution so that the wells have overflowed in the solution, and verify that the solution is evenly leveled with the gel on both sides;
6. Load 40.0 µL of the ladder in Lane-1;
7. Load 40.0 µL of each amplicon sample into the other wells;
8. Run the gel electrophoresis apparatus for 50 min by applying a voltage set to 106 V on the power source;
9. Remove the agarose gel from the gel electrophoresis apparatus and view it under UV light to visualize the bands (**Figure 4**);
10. Excise the amplicon gel band of interest from the agarose gel using a sterile scalpel blade;
11. Transfer the excised gel band (purified DNA amplicon) into a sterile 1.5-mL Eppendorf tube;
12. Store the purified DNA amplicon at 4 °C, which will be used in the DNA gel extraction procedure.

**Figure 4.**
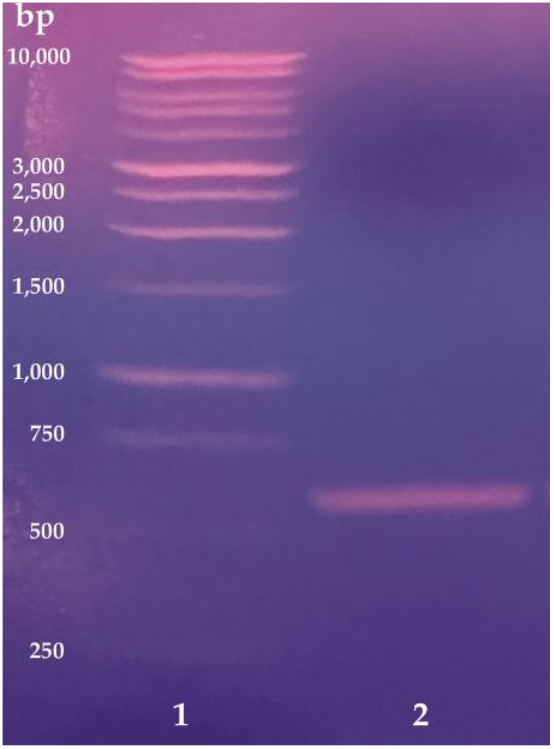
Purification of a truncated *Ao*COI gene amplicon (578 bp) by agarose gel electrophoresis. The *Ao*COI amplification by PCR involved *A. ornata* genomic DNA from one worm specimen and *Ao*COI-specific primers. Lane **1** is a 1-kb DNA ladder and lane **2** is a single sample of PCR amplicon product. Electrophoresis was performed using a 1.2% (w/v) agarose gel containing ethidium bromide with a 40 μL sample loading volume.

#### 3.2.4 Amplicon DNA Gel Extraction

1. Obtain the QIAquick Gel Extraction Kit (Qiagen);
2. Obtain the 1.5-mL Eppendorf tube containing the excised agarose gel band of the purified DNA amplicon of interest (see step 12 in section 3.2.3);
3. Add 1,200 µL of buffer QG (Qiagen);
4. Vortex and hand warm until the gel becomes fully dissolved;
5. Transfer the sample into a sterile 15 mL conical tube;
6. Add 500 µL of 100% isopropanol;
7. Mix gently with a micropipette;
8. Transfer 700 µL of the sample solution into a spin column;
9. Centrifuge at 13,000 rpm for 1.0 min;
10. Discard the flow-through;
11. Repeat steps 8 – 10 until there is no more sample solution left in the 15 mL conical tube;
12. Add 700 µL of buffer PE (Qiagen) into the spin column (note: 96% ethanol was added to the buffer PE concentrate provided in the kit);
13. Centrifuge at 13,000 rpm for 1.0 min;
14. Discard the flow-through;
15. Transfer the spin column into a sterile 1.5-mL Eppendorf tube;
16. Add 40.0 µL of buffer EB (Qiagen) directly onto the membrane of the spin column and allow to absorb for at least 1.0 min;
17. To elute the purified amplicon DNA, centrifuge at 13,000 rpm for 1.0 min;
18. Recover the flow-through by transferring it into a new sterile 1.5-mL Eppendorf tube that is properly labeled;
19. Store the purified gel-extracted DNA amplicon sample at -20 °C.

#### 3.2.5 Sanger DNA Sequencing

1. Obtain the purified gel-extracted DNA amplicon sample in buffer EB (see step 19 in section 3.2.4);
2. Thaw to room temperature from -20 °C;
3. Obtain an 8-strip of 0.2 mL PCR tubes with caps:
  a. In tube-1, perform the following;
    i. Add 10.0 μL of the purified DNA amplicon sample;
    ii. Add 2.5 μL of buffer EB (Qiagen);
    iii. Add 2.5 uL of 10.0 μM forward primer (*Ao*COI-FORW);
  b. In tube-2, perform the following;
    i. Add 10.0 μL of the purified DNA amplicon sample;
    ii. Add 2.5 μL of buffer EB (Qiagen);
    iii. Add 2.5 uL of 10.0 μM reverse primer (*Ao*COI-REVR);
4. Mail samples out to Azenta Life Sciences (Research Triangle Park, NC, USA) for Sanger DNA sequencing service at ambient temperature;
  a. DNA type: purified PCR product.
5. Analyze the DNA sequences by running a standard nucleotide BLAST search on the National Center for Biotechnology Information (NCBI) server against GenBank accession no. OQ322956.1 (**Supplementary Information**) using the default output parameters at https://blast.ncbi.nlm.nih.gov/.

### 3.3. Protein Purification

#### 3.3.1 Extraction and Purification of AoDHP_NS_ from A. ornata

1. Extraction of *Ao*DHP_NS_ from *A. ornata*;
  a. Followed a published protocol by Chen and colleagues [19] but introduced additional modifications;
  b. Chill a mortar and pestle at -80 °C;
  c. Select four *A. ornata* specimens (verified by DNA barcoding using the *Ao*COI gene; see section 3.2) that weigh out to approx. 5.0 g;
  d. Thaw specimens in a plastic weighing-boat from -80 °C to room temperature (**Figure 5a**);
  e. Add 10 mL of cold lysis buffer [100 mM potassium phosphate (pH 7.0)];
  f. Transfer all contents into the mortar (note: worms will re-freeze once they are placed into the mortar);
  g. Crush and grind the specimens with a pestle;
  h. Transfer liquid sample into a 50-mL conical tube;
  i. Continue to crush and grind the sample until a consistency of ground tissue is observed (**Figure 5b**);
  j. Transfer all sample into the 50-mL conical tube;
  k. Add an additional 10 mL of cold lysis buffer;
  l. Stir the sample using a spatula;
  m. Sonicate the sample at 0 °C (on ice) using an ultrasonic homogenizer, such as the QSonica Sonicator Q55 (Fisher Scientific) with the following settings (note: perform a total of 3 sonication cycles);
    i. Set the amplitude to 70%;
    ii. Apply a pulse by using the thumb switch every 1 sec for 60 sec;
    iii. Allow for a 60-sec rest period;
    iv. Stir lysed cells/broken tissue with a spatula;
    v. Repeat steps two more times;
  n. Set four layers of cheesecloth in a powder funnel;
  o. Place the funnel over a new 50-mL conical tube;
  p. Immediately pass the cell lysate through the cheesecloth (note: wear clean nitrile gloves);
  q. Create a protease inhibitor solution by dissolving one EDTA-free protease inhibitor tablet (Pierce) into 5 mL of lysis buffer;
  r. Add the protease inhibitor solution into the cell lysate;
  s. Add two heaping spatula-tip full amounts of potassium ferricyanide (solid) in order to oxidize *Ao*DHP_NS_ to the ferric form;
  t. Stir cell lysate until all of the potassium ferricyanide dissolves;
  u. Centrifuge the cell lysate at 11,400 rpm for 45 min at 4 °C using the Allegra X-30R refrigerated centrifuge (Beckman-Coulter) to pellet cellular debris;
  v. Recover the supernatant, which will be clarified from cellular debris, by transferring into a new 50-mL conical tube (**Figure 5c**);
    i. The sample will now be referred to as the *Ao*DHP_NS_ sample;
  w. Transfer the 25 mL *Ao*DHP_NS_ sample into a 6 – 8 kDa molecular weight cut-off dialysis membrane;
2. Purification: Dialysis;
  a. Run dialysis on the *Ao*DHP_NS_ sample (note: a total of 3 dialyses should be performed), as follows;
    i. Prepare 4 L of dialysis buffer [20 mM Tris (pH 7.4)] in a 4-L plastic beaker having large stir bar;
    ii. Set the beaker into a scientific refrigerator (temperature of 4°C) on a stir plate;
    iii. Place the dialysis membrane containing *Ao*DHP_NS_ into the dialysis buffer;
    iv. Dialyze for 4 hrs with moderate stirring of the dialysis buffer;
    v. Repeat steps two more times;
  b. Recover the *Ao*DHP_NS_ sample from the dialysis membrane;
  c. Centrifuge at 11,400 rpm for 45 min at 4 °C using the Allegra X-30R refrigerated centrifuge (Beckman-Coulter) to pellet any cellular debris;
  d. Recover the supernatant;
  e. Syringe filter the sample through a hydrophilic 0.22 μm PVDF filter (Millipore).
3. Purification: CM Sepharose Fast Flow Chromatography;
  a. Run CM Sepharose FF chromatography using an ÄKTA Prime Plus FPLC system (GE Healthcare), as follows;
  b. Prepare 1 L of filtered mobile phase A (MP-A) [20 mM Tris (pH 7.4)];
  c. Prepare 1 L of filtered MP-B [100 mM potassium phosphate (pH 7.0), 400 mM potassium chloride];
  d. Pack a column with CM Sepharose FF resin (GE Healthcare) [1.5 cm I.D. x 9 cm resin height];
  e. Pass 10 column volumes (CVs) of deionized water by gravity flow;
  f. Pass 10 CVs of MP-A (*vide supra*) by gravity flow;
  g. Connect the column to the FPLC;
  h. Start flowing MP-A at a flow-rate of 2.0 mL/min;
  i. Apply the following manual FPLC settings;
    i. Flow-rate = 2.0 mL/min;
    ii. Fraction size = 5.0 mL;
    iii. Pressure limit = 0.30 MPa;
    iv. Wavelength on UV detector = 280 nm;
  j. Inject the approximate 25 mL *Ao*DHP_NS_ sample in: 20 mM Tris (pH 7.4) onto the CM Sepharose FF column by using an ÄKTA 50-mL superloop (GE Healthcare);
  k. After injection is complete, run MP-A until a baseline is reached on the UV detector;
  l. Set a gradient elution to a final target of 100% MP-B with a gradient length of 70.0 mL;
  m. *Ao*DHP_NS_ elutes prior to the gradient elution and appears to the eye as a red-colored sample, and the sample is dissolved in MP-A;
  n. Pool the *Ao*DHP_NS_ fractions from CM Sepharose FF chromatography;
  o. Concentrate the sample to approximately 20 mL using an Amicon Ultra-15 centrifugal concentrator (MWCO = 10 kDa) by centrifuging at 4,100 rpm at 4 °C for 22 min;
4. Purification: Q-HP Chromatography;
  a. Run Q-HP chromatography using an ÄKTA Prime Plus FPLC system (GE Healthcare), as follows;
  b. Connect a pre-packed 5-mL Q-HP chromatography column (Cytiva) to the FPLC;
  c. Pass 10 CVs of deionized water at a flow-rate of 2.0 mL/min;
  d. Pass 10 CVs of MP-A [20 mM Tris (pH 7.4)] at a flow-rate of 2.0 mL/min;
  e. Continue flowing MP-A;
  f. Apply the following manual FPLC settings;
    i. Flow-rate = 2.0 mL/min;
    ii. Fraction size = 3.0 mL;
    iii. Pressure limit = 0.30 MPa;
    iv. Wavelength on UV detector = 280 nm;
  g. Inject the 20 mL of pooled and concentrated *Ao*DHP_NS_ sample in: MP-A that was recovered from the CM Sepharose FF chromatography (step 29n) onto the Q-HP column by using an ÄKTA 50-mL superloop (GE Healthcare);
  h. After injection is complete, run MP-A until a baseline is reached on the UV detector;
  i. Set a gradient elution to a final target of 100% MP-B [100 mM potassium phosphate (pH 7.0), 400 mM potassium chloride] with a gradient length of 100.0 mL;
5. Purification: Protein Purity Assessment by SDS-PAGE;
  a. Perform SDS-PAGE electrophoresis on all Q-HP fractions of *Ao*DHP_NS_ that have a red color to the eye;
  b. Apply the following SDS-PAGE conditions, as follows:
  c. Use a Mini-Protean TGX 4 – 15% polyacrylamide gel, 15-well, 15 μL volume capacity per well (Bio-Rad);
  d. Prepare a Laemmli (2x) (Bio-Rad)/β-mercaptoethanol solution as mixing 950 μL:5.0 μL (190:1 v/v), respectively;
  e. In separate 1.5-mL Eppendorf tubes, prepare each fraction with the Laemmli/β-mercaptoethanol solution as mixing 10 μL:10 μL (1:1 v/v), respectively;
  f. Denature SDS-PAGE samples by boiling in a water bath (100 °C) for 5 min;
  g. Allow samples to cool to room temperature;
  h. Pipet 12.0 μL of each sample per well;
  i. Pipet 12.0 μL of the Precision Plus Protein Dual Color Standards (Bio-Rad) for a ladder marker;
  j. Run the gel using an electrophoresis apparatus (Bio-Rad) with a Tris/glycine running buffer (1x) at 200 V for ∼25 – 30 min;
  k. Wash the gel with water;
  l. Stain the gel using a stain solution [0.1% (w/v) Coomassie R-250 in: 10% acetic acid, 40% methanol];
  m. Destain the gel using a destain solution [10% acetic acid, 40% methanol];
  n. Analyze the SDS-PAGE gel by visual inspection and determine the purest fractions of *Ao*DHP_NS_ that have a molecular weight of 15.5 kDa;
6. Purification: Buffer Exchange;
  a. Pool the purest fractions of *Ao*DHP_NS_;
  b. Buffer exchange *Ao*DHP_NS_ into assay buffer [100 mM potassium phosphate (pH 7.0)] by using a PD-10 desalting column (GE Healthcare), as follows:
  c. Concentrate the *Ao*DHP_NS_ sample to a volume of 2.5 mL using an Amicon Ultra-15 centrifugal concentrator (MWCO = 10 kDa) by centrifuging at 4,100 rpm at 4 °C for 22 min;
  d. Take a PD-10 desalting column and remove the cap above the column resin and drain the storage liquid (note: rinse this area six times with deionized water);
  e. Using scissors, cut off some plastic at the bottom of the column so that liquid can flow through;
  f. Mount the column on a ring stand and place a flask underneath the column to collect waste;
  g. Pass 25 mL of deionized water through the column;
  h. Pass 25 mL of assay buffer (*vide supra*) through the column.
  i. Under the bottom of the column, replace the flask with a collection
  j. tube (15 mL tube) and label it as “collection tube no. 1”;
  k. Add the 2.5 mL of *Ao*DHP_NS_ sample into the column;
  l. When exactly 2.0 mL of flow-through exits the column, switch to a second collection tube labeled as “collection tube no. 2”;
  m. Once the entire sample penetrates into the column bed, add 3.0 mL of assay buffer (*vide supra*);
  n. Collect all of the flow-through (red-colored protein) in collection tube no. 2 until the flow stops (note: the *Ao*DHP_NS_ sample that is buffer-exchanged into the new assay buffer is in this tube);
7. Purification: Concentration Determination and Adjustment;
  a. Perform a protein concentration determination and adjustment, as follows;
  b. Blank an Agilent 8453 UV-visible spectrophotometer using a 1.0 cm pathlength quartz cuvette microcell (operation vol. of 150 μL) (Starna Cells) filled with assay buffer [100 mM potassium phosphate (pH 7.0)];
  c. Repeat blanking procedure twice;
  d. Scan 150 μL of sample (*Ao*DHP_NS_ in: assay buffer);
  e. Observe the absorbance of the heme Soret band (406 – 418 nm);
  f. Observe the absorbance of the protein band (280 nm);
  g. Determine the Reinheitszahl (R_Z_) value, which is the A_Soret_/A_280_ (note: typical R_Z_ values for this preparation are in the range of 1.60 – 2.00);
  h. Use the Beer-Lambert Law (A = ε C l) to determine the concentration of the *Ao*DHP_NS_ sample (note: ε406nm = 116,400 M-1 cm-1 [26]);
  i. Adjust the concentration of *Ao*DHP_NS_ to 8.2 μM using assay buffer as a diluent.

**Figure 5.**
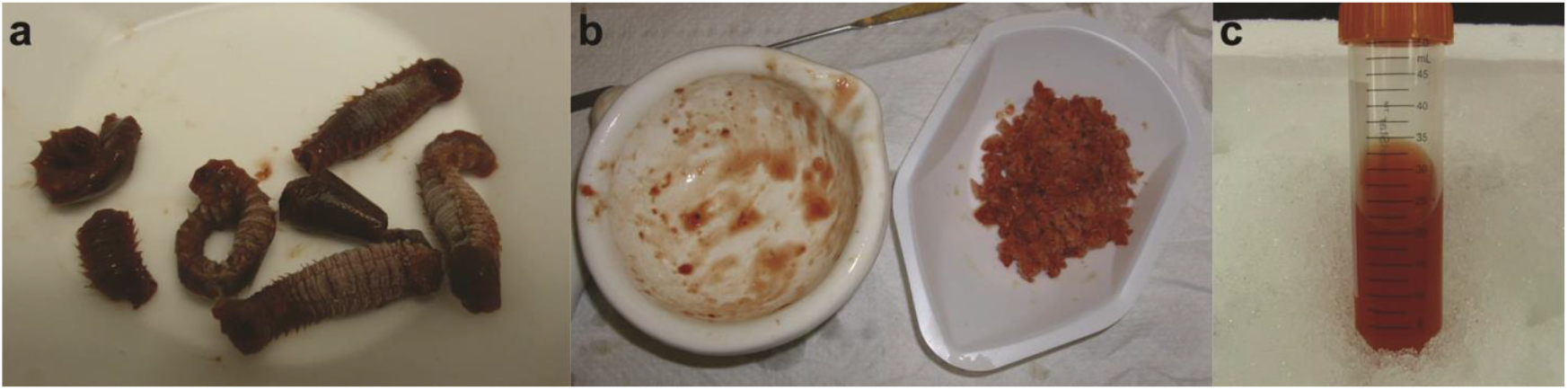
Extraction of *Ao*DHP_NS_ from *A. ornata* whole worm tissue. (a) *A. ornata* marine worms undergoing a thawing procedure from -80 °C to room temperature in a weighing-boat. (b) Ground worm tissue. (c) Impure *Ao*DHP_NS_ recovered in a 50-mL conical tube after being thoroughly sonicated, passed through four layers of cheesecloth, and centrifuged to remove cellular debris.

#### 3.3.2 Cloning, Overexpression, and Purification of Recombinant AoDHP_REC_-A

1. Cloning: Gene Synthesis;
  a. Through a gene synthesis service (Azenta Life Sciences, South Plainfield, NJ, USA), acquire plasmid DNA containing the gene for *A. ornata* dehaloperoxidase isoenzyme A, GenBank accession no. AAF97245.1 (**Supplementary Information**), cloned into a kanamycin-resistant pET-28a(+) *E. coli* expression vector;
    i. The gene should be cloned at the 5’ NcoI and 3’ HindIII restriction sites.
    ii. Codons for the gene synthesis should be optimized for protein expression in *E. coli*.
    iii. The construct should encode the 8-residue segment (bearing a 6x histidine tag): MGHHHHHH to precede the second residue, Gly-2; in addition, at the C-terminus of the protein, the final residue is Lys-138 (K138);
    iv. The plasmid is designated henceforth as pET-*Ao*DHP_REC_-A;
2. Overexpression: Transformation;
  a. All steps require the sterilization techniques used in molecular biology (e.g., sterile pipet tips, sterile 1.5-mL Eppendorf tubes, etc.);
  b. Transform plasmid pET-*Ao*DHP_REC_-A by the heat shock method into *E. coli* strain BL21(DE3) (New England Biolabs, Ipswich, MA);
  c. Thaw out a 50 μL aliquot of BL21(DE3) cells on ice for 15 min from -80 °C;
  d. Thaw plasmid DNA on ice for 15 min from -20 °C;
  e. Mix pET-*Ao*DHP_REC_-A (2.0 μL) with BL21(DE3) cells (50 μL) and allow the solution to incubate on ice for an additional 15 min;
  f. Place two LB – kanamycin plates in an incubator at 37 °C to warm up;
  g. Heat shock the cells by placing the tube containing the BL21(DE3)/pET*Ao*DHP_REC_-A mixture in a water bath set to 42 °C for 2.0 min;
  h. Immediately place the tube back on ice;
  i. Add 400 μL of LB broth;
  j. Incubate cells at 37 °C and shake at 250 rpm for 1.0 hr;
  k. Spread cells on the two LB – kanamycin plates by first pipetting 125 μL on one plate and 325 μL on the other plate (note: use a glass spreader that is sterilized by 91% isopropyl alcohol and flame);
  l. Incubate cells agar side up for 16 hrs (overnight);
  m. Observe colony growth on both plates to a good size and distribution;
3. Overexpression: 5-mL Starter Cultures;
  a. In a sterile 50-mL conical tube, mix LB broth (40 mL) with 50 mg/mL kanamycin (40.0 μL), and vortex the solution;
  b. Transfer 5 mL of LB/kan solution into seven culture tubes;
  c. Inoculate each tube containing LB/kan solution with a single colony of BL21(DE3)/pET-*Ao*DHP_REC_-A;
  d. Incubate starter cultures at 37 °C and shake at 220 rpm for 16 hrs;
4. Overexpression: 1000-mL Upscaled Cultures;
  a. Prepare 6x 1000-mL of 2xYT broth by mixing 31.0 g of 2xYT media and 1000 mL of steam-distilled water into separate 2-L flasks with baffles, add a cheesecloth lid with a rubber band, and add an aluminum foil cap with autoclave tape to hold it in place;
  b. Autoclave the six 1000-mL media and allow to cool down to room temperature;
  c. Perform the following steps per flask;
    i. Add 1000 μL of 50 mg/mL kanamycin and swirl to mix to homogeneity;
    ii. Inoculate with 5 mL of starter culture (from the 5-mL starter cultures step, *vide supra*);
    iii. Add 1000 μL of 5.0 mg/mL hemin (from porcine, Sigma) in: 0.1 M NaOH (sterile solution) and immediately swirl broth (note: the final pH of the 1 L broth should be checked by pH paper and it should have a pH of ∼7);
  d. Incubate the 1000-mL upscaled cultures at 37 °C and shake at 220 rpm for 14 – 17 hrs;
  e. Pellet cells by using a centrifuge;
    i. Spin media at 10 min at a time at 4 °C using a swinging bucket rotor in order to group cell mass in centrifugation bottles;
  f. Transfer cells into a hexagonal weigh boat using a large spatula;
  g. Observe a dark brownish pink cell pellet having a mass of 40 g;
  h. Cover the pellet with clear plastic wrap and label;
  i. Store cell pellet immediately at -80 °C;
5. Purification: Cell Lysis (Part 1);
  a. Thaw the 40 g BL21(DE3) cell pellet from -80 °C to room temperature;
  b. Place the cell pellet while it is frozen into a plastic beaker;
  c. Resuspend cells in 80 mL of cell lysis buffer [50 mM HEPES (pH 7.0), 300 mM NaCl] at room temperature;
  d. Add 0.5 g of lyophilized lysozyme into a 50-mL conical tube containing 10 mL of cell lysis buffer and swirl to fully dissolve;
  e. Add the lysozyme solution into the cell suspension;
  f. Stir cell suspension for 40 min at 4 °C;
  g. After the first 10 min passes during the stirring step, add two protease inhibitor tablets (EDTA-free);
  h. Sonicate the cell suspension for 30 min using a water bath sonicator at 0 °C (ice in the water of the water bath);
  i. Store lysed cell suspension at -20 °C overnight;
6. Purification: Cell Lysis (Part 2);
  a. Thaw cell lysate from -80 °C to room temperature for ∼2 hrs;
  b. Hand stir until all ice breaks apart;
    i. Observation: very high viscosity;
  c. Add 500 μL of 10 mg/mL DNase I;
  d. Add 500 μL of 16 mg/mL RNase A;
  e. Hand stir using a spatula for 3 min;
  f. Stir in a refrigerator (4 °C) for 1.0 hr;
    i. Observation no. 1: fully diminished viscosity (note: similar to the viscosity of milk);
    ii. Observation no. 2: color was a brownish pink;
  g. Store the cell lysate at -80 °C overnight;
7. Purification: Co-NTA Chromatography;
  a. Prepare 1.0 L of mobile phase A [50 mM HEPES (pH 7.0), 300 mM NaCl];
  b. Prepare 1.0 L of mobile phase B [50 mM HEPES (pH 7.0), 300 mM NaCl, 150 mM imidazole];
  c. Thaw the cell lysate from -80 °C to room temperature for -3 hrs;
    i. Observation: a low viscosity similar to that of milk);
  d. Centrifuge the cell lysate for 45 min at 11,400 rpm at 4 °C using a Beckman Coulter fixed-angle rotor;
  e. Collect the supernatant immediately and transfer into 50-mL conical tubes (note: the cell pellet debris in each bullet tube can be discarded since the protein is found in the supernatant);
  f. Pack a chromatography column with Talon Co2+-NTA resin with dimensions [1.5 cm (inside diameter) x 2.5 cm (height)];
  g. Connect the column to a P-1 peristaltic pump (GE Healthcare) and run the pump at room temperature;
  h. Flow 10 column volumes of deionized water through the column at a flow-rate of 2.0 mL/min;
  i. Flow 10 column volumes of mobile phase A (*vide supra*) at a flow-rate of 2.0 mL/min;
  j. Load *Ao*DHP_REC_-A sample (supernatant) onto the Co2+-NTA resin bed at a flow-rate of 2.0 mL/min;
  k. Disconnect the column from the P-1 peristaltic pump;
  l. Connect the column to an ÄKTA Prime Plus FPLC system (GE Healthcare) which is housed within a refrigerator at 4 °C;
  m. Turn on the UV lamp (λ = 280 nm);
  n. Start the fraction collector with a fraction volume set to 5.0 mL/fraction;
  o. Set a flow-rate of 2.0 mL/min;
  p. Flow mobile phase A (*vide supra*) until a baseline is reached;
  q. Perform an isocratic step up to 13% mobile phase B and continue to flow until an impurity peak is observed to elute (∼20 – 30 min);
  r. Perform an isocratic step up to 100% mobile phase B and continue to flow until all protein elutes;
  s. End the run on the FPLC;
  t. Pool fractions from the 100% mobile phase B elution step (vol. of ∼20 mL) into a 50-mL conical tube that proceeds to the next step;
  u. Store the pooled fraction set (*Ao*DHP_REC_-A) at 4 °C;
8. Purification: Dialysis;
  a. Oxidize the *Ao*DHP_REC_-A sample by addition of slight excess potassium ferricyanide;
  b. Prepare three 4-L solutions of dialysis buffer [10.0 mM potassium phosphate (pH 7.0)];
  c. Soak a dialysis membrane tubing (6 – 8 kDa MWCO) in distilled water for 5 min;
  d. Tie a knot on end of the dialysis tubing followed by filling the tubing with the 20 mL sample of *Ao*DHP_REC_-A;
  e. Tie a second knot on the other end of the tubing;
  f. Run the first dialysis session with stirring for 4 hrs at 4 °C;
    i. Observation: after 4 hrs has passed, the buffer in the beaker should have a yellowish tint to it;
  g. Run the second dialysis session with stirring for 4 hrs at 4 °C;
    i. Observation: after 4 hrs has passed, the buffer in the beaker should be colorless;
  h. Run the third dialysis session with stirring for 4 hrs at 4 °C;
    i. Observation-1: after 4 hrs has passed, the buffer in the beaker should be colorless;
    ii. Observation-2: the *Ao*DHP_REC_-A sample within the dialysis tubing should have minimal to no precipitation;
  i. Pull out the dialysis sample by wearing nitrile gloves and rinse it with deionized water;
  j. Place a powder funnel over a 50-mL conical tube;
  k. Recover the dialysate by carefully piercing the dialysis tubing with a cleaned razor blade above the funnel, and allow the *Ao*DHP_REC_-A sample to flow into the 50-mL conical tube;
    i. Observation: *Ao*DHP_REC_-A is in the Ferric oxidization state;
9. Purification: CM Sepharose Fast Flow Chromatography;
  a. Prepare 1.0 L of mobile phase A [10.0 mM potassium phosphate (pH 7.0)];
  b. Prepare 1.0 L of mobile phase B [38.0 mM potassium phosphate (pH 7.0)];
  c. Pack a chromatography column with CM Sepharose Fast Flow resin (GE Healthcare) with dimensions [1.5 cm (inside diameter) x 3.5 cm (height)];
  d. Connect the column to an ÄKTA Prime Plus FPLC system (GE Healthcare) (Temp. = 4 °C);
  e. Turn on the UV lamp (λ = 280 nm);
  f. Start the fraction collector with a fraction volume set to 3.0 mL/fraction;
  g. Set a flow-rate of 2.0 mL/min;
  h. Flow deionized water until 10 column volumes pass through;
  i. Flow mobile phase A (*vide supra*) until 10 column volumes pass through;
  j. Add the 20 mL of *Ao*DHP_REC_-A into an ÄKTA 50-mL superloop (GE Healthcare) and inject sample onto the column;
  k. After sample loads onto the column, disconnect the superloop;
  l. Flow mobile phase A (*vide supra*) until a baseline is reached;
  m. Perform a gradient elution from 0 – 100% mobile phase B with a 40.0 mL target volume;
  n. Continue to flow 100% mobile phase B until all protein elutes;
  o. End the run on the FPLC;
  p. Pool fractions all fractions that eluted from the gradient elution into a 50-mL conical tube;
  q. Store the pooled fraction set (*Ao*DHP_REC_-A) at 4 °C;
10. Purification: Buffer Exchanging Purified *Ao*DHP_REC_-A into Assay Buffer;
  a. Concentrate the *Ao*DHP_REC_-A pooled fraction set from CM Sepharose FF purification down to 1.0 mL using an Amicon Ultra-15 centrifugal concentrator (MWCO = 10 kDa) by centrifugation for 22 min at 4 °C at 4,100 rpm;
  b. The sample was diluted to 15 mL using assay buffer [100 mM potassium phosphate (pH 7.0)];
  c. Repeat the previous two steps, two more times;
  d. Concentrate the sample down to 1.5 mL using the Amicon Ultra-15 concentrator;
  e. Sample: purified *Ao*DHP_REC_-A in: 100 mM potassium phosphate (pH 7.0);
11. Purification: Protein Purity Assessment by SDS-PAGE;
  a. Perform SDS-PAGE electrophoresis on the *Ao*DHP_REC_-A fractions from all chromatography performed;
  b. Apply the following SDS-PAGE conditions, as follows:
  c. Use a Mini-Protean TGX 4 – 15% polyacrylamide gel, 15-well, 15 μL volume capacity per well (Bio-Rad);
  d. Prepare a Laemmli (2x) (Bio-Rad)/2-mercaptoethanol solution as mixing 950 μL:5.0 μL (190:1 v/v), respectively;
  e. In separate 1.5-mL Eppendorf tubes, prepare each fraction with the Laemmli/β-mercaptoethanol solution as mixing 10 μL:10 μL (1:1 *v/v*), respectively;
  f. Denature SDS-PAGE samples by boiling in a water bath (100 °C) for 5 min;
  g. Allow samples to cool to room temperature;
  h. Pipet 12.0 μL of each sample per well;
  i. Pipet 12.0 μL of the Precision Plus Protein Dual Color Standards (Bio-Rad) for a ladder marker;
  j. Run the gel using an electrophoresis apparatus (Bio-Rad) with a Tris/glycine running buffer (1x) at 200 V for ∼25 – 30 min;
  k. Wash the gel with water;
  l. Stain the gel using a stain solution [0.1% (w/v) Coomassie Brilliant Blue R-250 in: 10% acetic acid, 40% methanol];
  m. Destain the gel using a destain solution [10% acetic acid, 40% methanol];
  n. Analyze the SDS-PAGE gel by visual inspection and determine the purest fractions of *Ao*DHP_REC_-A (note: with the 8-residue segment MGHHHHHH that precedes Gly-2, the molecular weight of *Ao*DHP_REC_-A is 16,540 Da);
12. Purification: Concentration Determination and Adjustment;
  a. Turn on the UV and visible lamps on an Agilent 8453 spectrophotometer and allow to warm up for at least 10 min;
  b. Observe the UV-visible spectrum from a wavelength range of 200 – 800 nm;
  c. Blank the instrument three times using 100 mM potassium phosphate (pH 7.0) in a 1.0 cm pathlength quartz microcuvette (sample vol. of 150 μL);
  d. Scan the blank;
  e. Mix *Ao*DHP_REC_-A solution with 100 mM potassium phosphate (pH 7.0) (30 μL : 120 μL);
  f. Scan the diluted sample and record the absorbance at a wavelength maximum at 407 nm.
  g. Solve for concentration using the Beer – Lambert Law (A = εCl), where ε407 = 116,400 M-1 cm-1 [26];
  h. Back-calculate by using the dilution equation C_i_V_i_ = C_f_V_f_ for the initial concentration;
  i. Adjust the concentration of the purified *Ao*DHP_REC_-A sample to be [protein] = 8.2 μM;

### 3.4 Trypsin Digestion LC-MS/MS

#### 3.4.1 Sample Preparation of AoDHP_NS_ and AoDHP_REC_-A for Trypsin Digestion

1. Perform an SDS-PAGE (similar to step 31 from section 3.3.1, *vide supra*) on purified samples of *Ao*DHP_NS_, *Ao*DHP_REC_-A (6x-His-tagged; control-1), and *Ao*DHP_REC, NHT_-A (non-His-tagged form; control-2) (**Figure 6**);
2. Cut out four gel bands of *Ao*DHP_NS_ using a clean razor blade;
3. Cut out four gel bands of non-His-tagged *Ao*DHP_REC_-A (*Ao*DHP_REC, NHT_-A) using a razor blade.
  a. Note: *Ao*DHP_REC, NHT_-A was provided to our laboratory by Prof. A. Reza Ghiladi (North Carolina State University, Dept. of Chemistry). The protein was overexpressed in *E. coli* and purified by traditional biochemical methods. The work followed a previously published protocol described by de Serrano and colleagues [27];
4. For each set of samples, mince the gel band pieces into 2.0 mm x 2.0 mm x 2.0 mm cubes (do not crush) and transfer into separate sterile 1.5-mL Eppendorf tubes (namely, one tube for *Ao*DHP_NS_ and one tube for *Ao*DHP_REC, NHT_-A);
5. Wash the gel band pieces twice, as follows:
  a. Cover the gel band pieces in 500 μL of the gel wash solution [acetonitrile: 50 mM Tris (pH 8.2) (1:1 *v/v*) and shake vigorously at 37 °C, 220 rpm for 20 min;
  b. Decant and discard the gel wash solution;
6. Dry the gel band pieces:
  a. Cover the gel band pieces in the Eppendorf tube with 40.0 μL of 100% acetonitrile (neat) and soak for 2.0 min at room temperature;
  b. Remove all acetonitrile from the tube;
7. Prepare a fresh stock solution of 0.1 μg/μL trypsin from 20 μg of trypsin;
  a. Dissolve 20 μg of trypsin (Promega, cat. No. V5111) into 200 μL of 50 mM Tris (pH 8.2) and store on ice;
8. Prepare a fresh working solution of 10 ng/μL trypsin;
  a. Mix 20 μL of 0.1 μg/μL trypsin with 180 μL of 50 mM Tris (pH 8.2);
9. Swell the gel band pieces for 30 min with 40 μL of 10 ng/μL trypsin at room temperature;
  a. Note: the gel band pieces were fully soaked in the trypsin solution, where 40 μL was a good lower limit volume;
  b. Check the gel band pieces after 30 min to verify that the trypsin solution was still soaking the gels;
    i. Added 20 μL of 10 ng/μL trypsin to the 1.5-mL Eppendorf tube of the gel band pieces to make the total volume 60 μL;
    ii. Incubate the sample at 37 °C for 16-18 hrs;
10. Centrifuge the sample at 13,000 rpm for 1.0 min;
  a. Recover the supernatant and place into a separate 1.5-mL Eppendorf tube (label as: “Initial Supernatant;”
  b. Transfer the gel band pieces into a new 1.5-mL Eppendorf tube;
11. Extract the peptide fragments;
  a. Prepare an extraction solution [1% (v/v) acetic acid, 2% (v/v) acetonitrile];
  b. Add 120 μL of the extraction solution to the gel band pieces so that they become submerged;
  c. Sonicate for 5.0 min at room temperature using a water-bath sonicator;
  d. Incubate at room temperature for 30 min and vortex at every 10 min;
  e. Centrifuge at 13,000 rpm for 1.0 min;
  f. Recover the supernatant solution surrounding the gel band pieces;
  g. Add the “Initial Supernatant” solution to this recovered supernatant solution;

**Figure 6.**
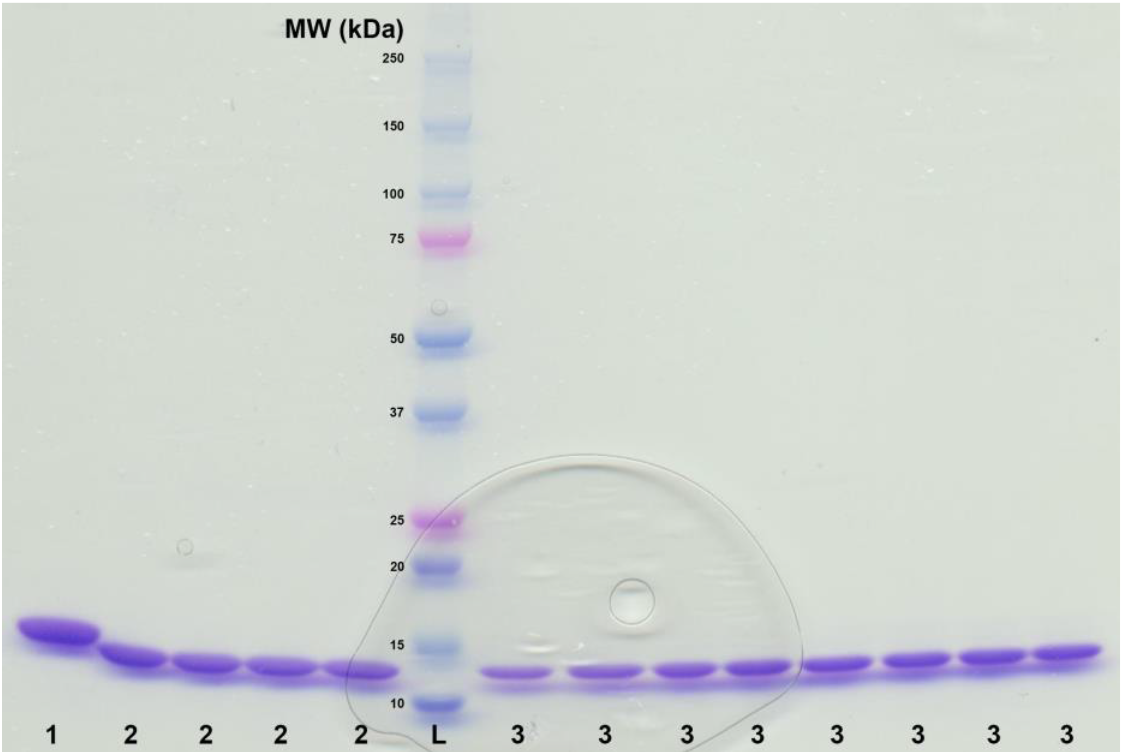
SDS-PAGE of sample **1** representing 6x-His-tagged *Ao*DHP_REC_-A (control-1; MW of 16,540 Da); sample **2** representing non-His-tagged *Ao*DHP_REC, NHT_-A (control-2; MW of 15,660 Da); and sample **3**: *Ao*DHP_NS_ (approx. 15.7 kDa). Sample **L** represents the molecular weight marker ranging from 10 – 250 kDa (Precision Plus Protein Dual Color Standards; Bio-Rad). The 15 kDa standard (within **L**) exhibited slightly different electrophoretic mobility compared to samples **2** and **3**, which have similar molecular masses. For trypsin digestion, four bands of sample **2** and four bands of sample **3** were cut from the gel using a cleaned razor blade. Electrophoresis was performed using a 4 – 15% polyacrylamide gel that was stained with Coomassie Brilliant Blue R-250 with a 12 μL sample loading volume per well.

#### 3.4.2 LC-MS/MS of Trypsin Digested Samples

a. Analyze tryptic digests by LC-MS/MS using an Easy NanoLC 1000 (Thermo Scientific, USA) coupled to an Orbitrap Elite mass spectrometry system (Thermo Scientific, USA);
b. Desalt and preconcentrate digests onto a 2 cm x 100 µm I.D. Pepmap C18 (5 µm particle) trapping column (Thermo Scientific, USA), and then elute onto and separate using an in-house packed PicoFrit (New Objective, USA) 75 µm I.D. x 25 cm Magic C18 column (3 µm particle) (Bruker Scientific) with a 30 min linear gradient from 2% mobile phase B to 40% mobile phase B (A = 2% acetonitrile in water, 0.1% formic acid; B = acetonitrile, 0.1% formic acid) at 300 nL/min;
c. An electrospray voltage of 2.8 kV was applied to the PicoFrit to ionize peptides in the nanoelectrospray ion source of the Obitrap Elite;
d. Data-dependent MS/MS data was collected utilizing the Orbitrap analyzer for both precursor and product ion mass analysis using a Top 5 method with higher-energy collisional activation (HCD) for product ion generation in the HCD cell;
e. A normalized collision energy of 27 V was used to induce precursor fragmentation;
f. Raw data files from the LC-MS/MS acquisitions were processed using Proteome Discoverer (Thermo Scientific, USA) and protein identifications were made by searching the SwissProt database using the MASCOT search engine (Matrix Science, UK) allowing for Met oxidation as a variable modification (**Figure 7**);

**Figure 7.**
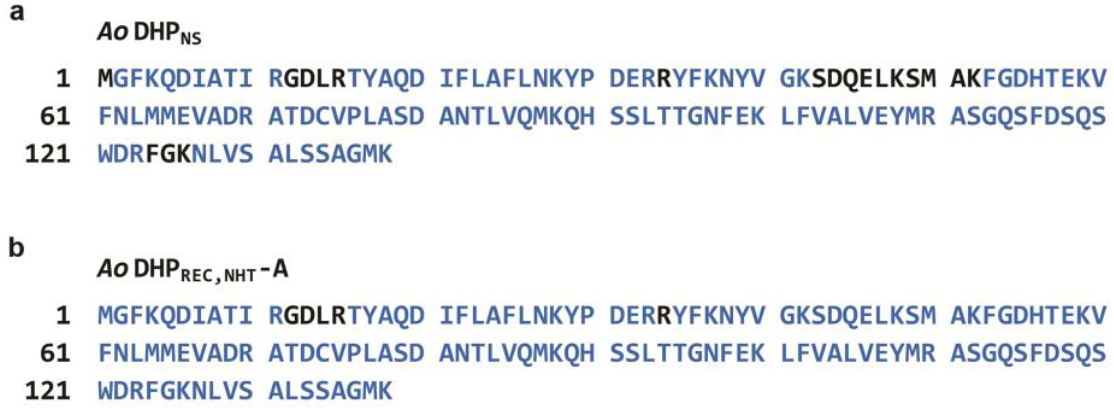
Analysis of trypsin digested samples for (a) *Ao*DHP_NS_ and (b) *Ao*DHP_REC, NHT_-A by LC-MS/MS. With respect to *A. ornata* dehaloperoxidase A (138 residues) (GenBank accession no. AAF97245.1), blue color-coding represents the matched peptides. Additionally, *Ao*DHP_NS_ and *Ao*DHP_REC, NHT_-A revealed protein sequence coverages of 86.2% and 96.4%, respectively. The proteolytic enzyme trypsin cleaves peptide bonds from the C-terminal direction of its target protein on lysine (K) or arginine (R) residues, unless they are next to a proline (P) residue.

#### 3.4.3 Peroxidase Assay of AoDHP_NS_ and AoDHP_REC_>-A

a. Perform a standard peroxidase assay (**Figure 8**), which is based on the oxidative dechlorination of 2,4,6-trichlorophenol (TCP) into 2,6-dichloro-1,4-benzoquinone (DCQ) when *Ao*DHP is exposed to hydrogen peroxide [14];
b. Note: in this section, *Ao*DHP is a general term that may refer to either *Ao*DHP_NS_ or *Ao*DHP_REC_-A, which depends of the form that the analyst decides to test;
c. Thaw purified 8.2 μM *Ao*DHP in assay buffer [100 mM potassium phosphate (pH 7.0)] from -80 to 0 °C (on ice);
d. Prepare 1.0 mL of 165 mM H_2_O_2_ in assay buffer by mixing 16.9 μL of 9.79 M H_2_O_2_ (Sigma, cat. no. 216763-100ML) with 983.1 μL of assay buffer (store on ice); note: 30% wt.% H_2_O_2_ = 9.79 M H_2_O_2_;
e. Prepare 1.0 mL of 10.0 mM H_2_O_2_ in assay buffer by mixing 60.6 μL of 165 mM H_2_O_2_ with 939.4 μL of assay buffer (store on ice);
f. Prepare 2.0 mL of 500 μM H_2_O_2_ in assay buffer by mixing 100 μL of 10.0 mM H_2_O_2_ with 1,900 μL of assay buffer (store on ice);
g. Prepare 2.0 mL of 10.0 mM TCP in 100% DMSO;
h. Perform an *Ao*DHP assay with the following parameters: *N=3*; variable [H_2_O_2_]; constant [TCP]; 8 reactions; reaction volume of 200 μL; 90 sec reaction time (time optimum); and scanning 70 μL of sample using a 96-well, clear, F-bottom, UV-Star microplate (Greiner Bio-One) in a BMG VANTAStar F microplate reader on UV-visible absorbance mode; record UV absorbance readings at λmax = 245 nm, 276 nm, and 312 nm; a TCP standard curve was used to determine the concentration of TCP consumed in the assay;
i. Reactions were carried out in assay buffer containing 10% (*v/v*) DMSO, co-substrate ([H_2_O_2_] ranged from 3.125 – 100.0 μM), substrate (1.0 mM TCP), and initiated with dehaloperoxidase (0.85 μM *Ao*DHP);
j. Perform an *Ao*DHP assay with the following parameters: *N=3*; constant [H_2_O_2_]; variable [TCP]; 8 reactions; reaction volume of 200 μL; 90 sec reaction time (time optimum); and scanning 70 μL of sample using a 96-well, clear, F-bottom, UV-Star microplate in a BMG VANTAStar F microplate reader on UV-visible absorbance mode; record UV absorbance readings at λmax = 245 nm, 276 nm, and 312 nm; a TCP standard curve was used to determine the concentration of TCP consumed in the assay;
k. Reactions were carried out in assay buffer containing 10% (*v/v*) DMSO, co-substrate (500 μM H2O_2_), substrate ([TCP] ranged from 7.813 – 250.0 μM), and initiated with 1.0 μM *Ao*DHP;
l. Process data on *Excel* and *GraphPad Prism 6.0* to determine Michaelis-Menten plots and kinetics parameters (**Table 1 and Figure 9**);
m. Observe a close similarity in *Ao*DHP kinetics parameters between source organisms from a side-by-side comparison (**Table 1**).

**Figure 8.**
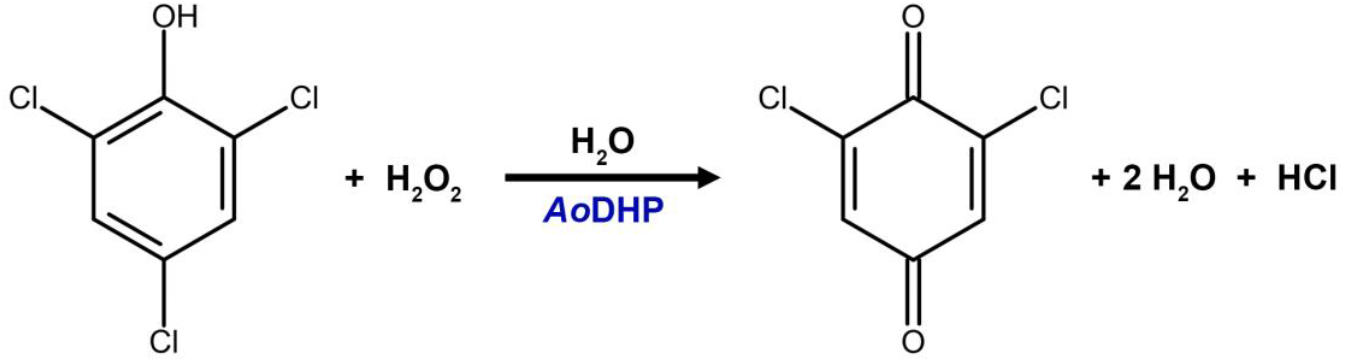
Peroxidase reaction for the oxidative dechlorination of 2,4,6-trichlorophenol into 2,6-dichloro-1,4-benzoquinone catalyzed by *Ao*DHP and hydrogen peroxide.

**Table 1.** Michaelis-Menten parameters for *Ao*DHP_NS_ and *Ao*DHP_REC_-A.^a^.

| Enzyme | Substrate | $K_M$ ($\mu\text{M}$) | $V_{\text{MAX}}$ ($\mu\text{M s}^{-1} \mu\text{g prot}^{-1}$) | $k_{\text{cat}}$ ($\text{s}^{-1} \mu\text{g prot}^{-1}$) | $k_{\text{cat}}/K_M$ ($\mu\text{M}^{-1} \text{s}^{-1} \mu\text{g prot}^{-1}$) |
| --- | --- | --- | --- | --- | --- |
| *AoDHP*<sub>NS</sub> | TCP | $51.2 \pm 24.6$ | $0.1095 \pm 0.0479$ | $0.0913 \pm 0.0399$ | $1.84 \times 10^{-3} \pm 2.24 \times 10^{-4}$ |
| | $\text{H}_2\text{O}_2$ | $2.58 \pm 2.08$ | $0.9148 \pm 0.2944$ | $1.289 \pm 0.4146$ | $8.01 \times 10^{-1} \pm 6.47 \times 10^{-1}$ |
| *AoDHP*<sub>REC-A</sub> | TCP | $11.0 \pm 11.2$ | $0.0618 \pm 0.0193$ | $0.0515 \pm 0.0160$ | $7.76 \times 10^{-3} \pm 5.40 \times 10^{-3}$ |
| | $\text{H}_2\text{O}_2$ | $5.67 \pm 5.66$ | $0.4966 \pm 0.1133$ | $0.5842 \pm 0.1332$ | $1.98 \times 10^{-1} \pm 1.82 \times 10^{-1}$ |

<sup>a</sup> Number of replicates, $N=3$ .

**Figure 9.**
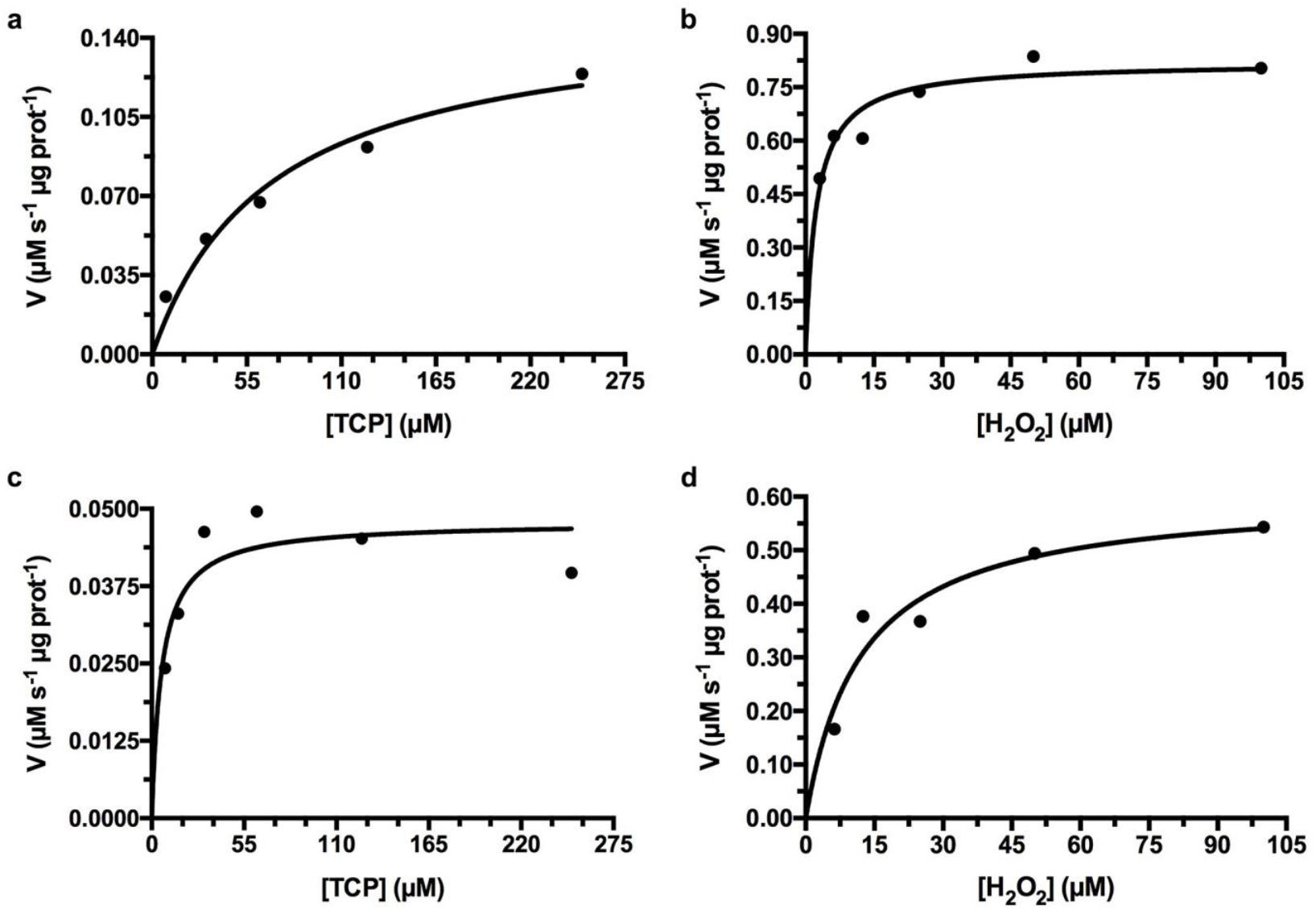
Enzyme kinetics of *Ao*DHP (natural source *vs*. recombinant) as represented by Michaelis-Menten plots for the peroxidase reaction. Panels (a and b) show rates of 2,6-dichloro-1,4-benzoquinone (DCQ) formation (90-sec time course) as a function of (a) [TCP] and (b) [H_2_O_2_] in the presence of *Ao*DHP_NS_. Panels (c and d) show rates of DCQ formation (90-sec time course) as a function of (c) [TCP] and (d) [H_2_O_2_] in the presence of *Ao*DHP_REC_-A. In these representative single experiments, the following Michaelis-Menten parameters were determined, as follows: in panel (a), *Ao*DHP_NS_ [K_M_ (TCP) = 69.2 μM, V^MAX^ (TCP) = 0.1519 μM s^-1^ μg prot^-1^, R^2^ = 0.9644] and in panel (b), *Ao*DHP_NS_ [K_M_ (H_2_O_2_) = 2.3 μM, V^MAX^ (H_2_O_2_) = 0.8198 μM s^-1^ μg prot^-1^, R^2^ = 0.9773]. In panel (c), *Ao*DHP_REC_-A [K_M_ (TCP) = 5.9 μM, V^MAX^ (TCP) = 0.04786 μM s^-1^ μg prot^-1^, R^2^ = 0.9262] and in panel (d), *Ao*DHP_REC_-A [K_M_ (H_2_O_2_) = 12.1 μM, V_MAX_ (H_2_O_2_) = 0.6057 μM s^-1^ μg prot^-1^, R^2^ = 0.9617].

## 4. Results and Discussion

There is currently a lack of discoveries made for other potential multifunctional catalytic hemoglobins originating from marine polychaetes. The outlined procedures herein will help support the accurate isolation, extraction, and characterization of hemoglobins from such macrobenthic infauna, since *A. ornata* was utilized as a representative model system. Notably, the methods have been designed for researchers that are not too familiar with the nuances involved in the visual taxonomy of polychaete worms, and the DNA barcoding approach will facilitate with accurate species identification.

In performing the molecular taxonomic species identification of *A. ornata*, we first began using the canonical universal COI primer by Folmer and colleagues, LCO1490 [5’-GGTCAACAAATCATAAAGATATTGG-3’] and HCO2198 [5’-TAAACTTCAGGGTGACCAAAAAATCA-3’], revealing only limited success [28,29]. In the PCR amplification reaction using *Taq* DNA polymerase, an amplicon product of interest (ca. 710-bp fragment) would result as was observed on a 1.2% agarose gel with ethidium bromide; however, typically a 30-bp fragment was only detected from DNA sequencing using either the forward or reverse primer. Furthermore, such results would only give rise to poor QS and CRL values of less than a value of 25 for each (for either the forward or reverse primers). Our approach was adjusted so that we could implement an *A. ornata*-specific primer set (*Ao*COI-FORW/*Ao*COI-REVR, see above in section 3.2.2.), which incidentally truncated our gene to a 578-bp fragment of the COI gene. On gel electrophoresis, the amplicon band appeared in the expected region (**Figure 4**), and more importantly, the DNA sequencing results showed an impressive improvement to the QS and CRL scores, particularly with the forward primer (typical scores of 42 for QS and 288 for CRL). With the forward primer, DNA sequencing revealed a sequence having 325 bp with matching identities to GenBank accession no. OQ322956.1 (**Supplementary Information**) from a nucleotide BLAST search on the NCBI server. We suspect that the specific *A. ornata*-COI primer set works better with the Sanger DNA sequencing reaction. The *Ao*COI-FORW and *Ao*COI-REVR primers were designed to have low occurrence of hairpin formation. If polychaete species are to be investigated for which there does not exist a COI gene in an online repository, and the LCO1490/HCO2198 primer set does not work appropriately, specific primers could be developed by modeling from known species having conservative sequences in the COI gene. The *Ao*COI primer set is most likely too specific for the DNA barcoding to be used in any other marine polychaetes unless substitution of a non-Watson-Crick nucleotide base is used, such as inosine. This replacement strategy may afford a primer with increased flexibility on binding to the genomic DNA.

On a comparison between the harvest of *Ao*DHP from *A. ornata* specimens vs. recombinant overexpression in *E. coli*, in both cases the protein yield was quite high. Isolation of *Ao*DHP_NS_ from *A. ornata* cannot resolve between the two isoenzymes, *Ao*DHP-A and *Ao*DHP-B; thus, a mixture of isoenzymes is most likely being purified in the sample.

Two key characterization methods, such as a trypsin digestion LC-MS/MS experiment and a TCP peroxidase assay were implemented in order to confirm the similarity of *Ao*DHP produced from two different source organisms, such as *A. ornata* marine worms and *E. coli* bacteria. In order to analyze the trypsin digested *Ao*DHP samples by LC-MS/MS, a brand-new trapping and analytical LC column was utilized. The only protein sample which was passed through this column prior to the *Ao*DHP samples was a tryptic digest of bovine serum albumin (BSA), which served as an in-house standard control sample for the purpose of system suitability. A volume of 10.0 μL was injected and analyzed for each of the samples, starting first with *Ao*DHP_NS_, followed by two BSA standard injections, and then, proceeding with the injection of *Ao*DHP_REC, NHT_-A. We made the decision to inject *Ao*DHP_NS_ onto the LC column first in order to avoid having any possible carryover of *Ao*DHP_REC, NHT_-A – unique, C-terminal peptides (analytical carryover) into a subsequent injection of *Ao*DHP_NS_. Using this approach eliminated the possibility of analytical carryover that would have otherwise confounded our interpretation of the results. *Ao*DHP_NS_ as compared to *Ao*DHP_REC, NHT_-A revealed a very high similarity based on protein sequence coverage obtained from both samples. **Figure 7a** shows that *Ao*DHP_NS_ has almost full coverage (86.2% matching to *Ao*DHP-A, GenBank accession no. AAF97245.1); whereas in **Figure 7b**, *Ao*DHP_REC, NHT_-A also shows almost full protein sequence coverage (96.4% matching to *Ao*DHP-A). Furthermore, protein sequence matching was extensive across the entire polypeptide chain of both *Ao*DHP samples (from the N-terminus to the C-terminus). Peroxidase activity exhibited by *Ao*DHP_NS_ and *Ao*DHP_REC_-A was evaluated, and Michaelis-Menten parameters were determined with respect to substrates TCP and H_2_O_2_ (**Table 1**). For *Ao*DHP_NS_, TCP had a K_M_ of 51.2 ± 24.6 μM and a VMAX of 0.1095 ± 0.0479 μM s-1 μg prot-1. H_2_O_2_ had a K_M_ of 2.58 ± 2.08 μM and a VMAX of 0.9148 ± 0.2944 μM s-1 μg prot-1. In the case of *Ao*DHP_REC_-A, TCP had a K_M_ of 11.0 ± 11.2 μM and a VMAX of 0.0618 ± 0.0193 μM s-1 μg prot-1. H_2_O_2_ had a K_M_ of 5.67 ± 5.66 μM and a VMAX of 0.4966 ± 0.1133 μM s-1 μg prot-1. The *Ao*DHP_NS_ or *Ao*DHP_REC_-A K_M_ values with respect to H_2_O_2_ that were determined in this study had slightly lower values to two previous reports involving *Ao*DHP_REC_-A with reported K_M_ values for H_2_O_2_ of 23 ± 1.2 μM [30] and 16 ± 1 μM [17]. Since it is possible that our collected *A. ornata* specimens bearing *Ao*DHP_NS_ potentially held a mixture of *Ao*DHP-A and *Ao*DHP-B, and regarding its K_M_ with respect to TCP (**Figure 9a** and **Table 1**), our observed value was also slightly lower in comparison to a previous report for that of *Ao*DHP_REC_-B and TCP (K_M_ of 210 ± 23 μM) [17]. We have determined that the kinetic parameters being observed between *Ao*DHP_NS_ and *Ao*DHP_REC_-A within this study revealed similar results. We also observed reproducibility from earlier studies, thus confirming our expectation that *Ao*DHP_NS_ was similar to *Ao*DHP_REC_-A.

The methods outlined herein efficiently describe the isolation, molecular taxonomy, extraction, and key confirmatory characterizations of hemoglobin (dehaloperoxidase) from *A. ornata* polychaetes that can be performed. Future studies should have a focus on field collections of other marine worm species using the isolation and molecular taxonomy methods presented. By especially doing so, more discoveries of novel enzymatic hemoglobins are predicted to follow.

## Author Contributions

Conceptualization, E.L.D.; data curation, A.L.H., V.R.S., H.V.N., L.A.P., R.K.B., J.L.S., S.A.B, and E.L.D.; formal analysis, A.L.H., V.R.S., H.V.N., L.A.P., J.L.S., S.A.B, and E.L.D.; funding acquisition, E.L.D. and V.R.S.; investigation, E.L.D.; resources, E.L.D.; software, E.L.D.; supervision, E.L.D.; validation, E.L.D.; visualization, E.L.D.; writing—original draft, A.L.H., V.R.S., and E.L.D.; and writing—review & editing, A.L.H., V.R.S., H.V.N., L.A.P., R.K.B., J.L.S., S.A.B, and E.L.D. All authors have read and agreed to the published version of the manuscript.

## Data Availability Statement

Data pertaining to key findings is contained within the article.

## Acknowledgments

The authors thank Prof. Reza A. Ghiladi at North Carolina State University, Dept. of Chemistry in Raleigh, North Carolina for providing purified *Ao*DHP_REC, NHT_-A that was used as a control in the trypsin digestion LC-MS/MS experiment. The authors also thank Lindsey R. Baker for technical support in the experimentation.

