## Supplementary Information for "The Multifunctional Catalytic Hemoglobin from *Amphitrite ornata*: Protocols on Isolation, Taxonomic Identification, Protein Extraction, Purification, and Characterization"

### Table of Contents

**I.** **Relevant GenBank Entries**

**a.** Nucleotide sequence for *Amphitrite ornata* mitochondrial cytochrome c oxidase subunit I (*Ao*COI) (GenBank accession no. OQ322956.1)….…………**S3**

**b.** Amino acid sequence for *Amphitrite ornata* dehaloperoxidase isoenzyme A (*Ao*DHP-A) (GenBank accession no. AAF97245.1)………...……**S4**

### I. Relevant GenBank Entries

#### a. Nucleotide sequence for *Amphitrite ornata* mitochondrial cytochrome c oxidase subunit I (*Ao*COI) (GenBank accession no. OQ322956.1)

*Amphitrite ornata* isolate SERCINVERT2482 cytochrome oxidase subunit 1 (COI) gene, partial cds; mitochondrial

GenBank: OQ322956.1

Number of Nucleotide Bases: 658

Source: https://www.ncbi.nlm.nih.gov/nuccore/OQ322956.1

FASTA Format

>OQ322956.1 *Amphitrite ornata* isolate SERCINVERT2482 cytochrome oxidase subunit 1 (COI) gene, partial cds; mitochondrial

```
AACTCTATATTTTATTTTTGGTATTTGAGGAGGCTTATTAGGAACCTCCATAAGATTACTAATTCGAA
TTGAACTTGGACAACCAGGAGCTTTTCTTGGAAGAGATCAACTGTATAACACAGTAGTAACAGCCCAT
GGTCTACTTATAATTTTTTTTCTAGTTATACCGATCCTTATTGGGGGATTCGGAAATTGATTACTACC
TCTTATATTAGGAGCACCCGATATAGCTTTCCCACGAATAAATAATATAAGATTTTGATTTTTACCTC
CTGCCCTTCTTCTATTACTTAGTTCAGCAGCTGTAGAAAAAGGTGTAGGTACAGGATGAACTGTGTAT
CCTCCTTTATCAAGAAATCTAGCACACGCTGGCCCCTCTGTAGATCTAGCTATTTTTTCCCTACATTT
AGCTGGGATCTCCTCAATCCTAGGAGCAATTAATTTTATTACAACAGTAGCTAACATACGATGAAAAG
GACTACGACTAGAACGAATCCCTCTATTTGTTTGAGCAGTTAATATTACTGTTATTTTACTTCTATTA
TCCTTACCAGTTCTAGCTGGTGCAATCACTATATTATTAACAGACCGTAATGTTAATACTTCTTTCTT
TGACCCATCAGGGGGAGGGGACCCAATTCTTTATCAACACTTATTT
```

**b. Amino acid sequence for *Amphitrite ornata* dehaloperoxidase isoenzyme A (*Ao*DHP-A) (GenBank accession no. AAF97245.1)**

*Amphitrite ornata* dehaloperoxidase A

GenBank: AAF97245.1

Number of Amino Acids: 138

Source: https://www.ncbi.nlm.nih.gov/protein/AAF97245.1

FASTA Format

>AAF97245.1 dehaloperoxidase A [*Amphitrite ornata*]

MGFKQDIATIRGDLRTYAQDIFLAFLNKYPDERRYFKNYVGKSDQELKSMAKFGDHTEKVFNLMMEVA

DRATDCVPLASDANTLVQMKQHSSLTTGNFEKLFVALVEYMRASGQSFDSQSWDRFGKNLVSALSSAG

MK
